# Transcriptome analysis reveals the difference between “healthy” and “common” aging and their connection with age-related diseases

**DOI:** 10.1101/747956

**Authors:** Lu Zeng, Jialiang Yang, Shouneng Peng, Jun Zhu, Bin Zhang, Yousin Suh, Zhidong Tu

## Abstract

A key goal of aging research is to understand mechanisms underlying healthy aging and use them to develop methods to promote the human healthspan. One approach is to identify gene regulations differentiating healthy aging from aging in the general population (i.e., “common” aging). In this study, we leveraged GTEx (Genotype-Tissue Expression) project data to investigate “healthy” and “common” aging in humans and their interconnection with diseases.

We selected GTEx donors who were not annotated with diseases to approximate a “healthy” aging cohort. We then compared the age-associated genes derived from this cohort with age-associated genes from our “common” aging cohort which included all GTEx donors; we also compared the “healthy” and “common” aging gene expressions with various disease-associated gene expression to elucidate the relationships among “healthy”, “common” aging and disease. Our analyses showed that 1. “healthy” and “common” aging shared a large number of gene regulations; 2. Despite the substantial commonality, “healthy” and “common” aging genes also showed distinct function enrichment, and “common” aging genes had a higher enrichment for disease genes; 3. Disease-associated gene regulations were overall different from aging gene regulations. However, for genes regulated by both, their regulation directions were largely consistent, implying some aging processes could increase the susceptibility to disease development; and 4. Possible protective mechanisms were associated with the “healthy” aging gene regulations.

In summary, our work highlights several unique features of human “healthy” aging program. This new knowledge can be used for the development of therapeutics to promote human healthspan.

## Introduction

Human life expectancy has increased by more than 30 years in the past 165 years in developed countries like the US (Christensen et al. 2009). By 2050, two billion of the estimated nine billion people on earth will be older than 60 (Burch et al. 2014). Aging is the major risk factor for many age-related diseases, including cancers, metabolic diseases, neurodegenerative diseases, and cardiovascular diseases (Johnson et al. 2015). The prevalence of age-related comorbidity is high, as over 80% of the elderly population having at least one chronic diseases (CDC 2011). Helping this 80% of the aging population to live with improved health condition has become a major task for human aging and geroscience research, which can have major impact to socioeconomics and humanity.

It remains elusive why the fortunate 20% of individuals above 65 years of age can live without any major health issues while the rest majority have to endure one or more chronic course of illness. Since very long-lived individuals (e.g., centenarians) tend to have a lower incidence of chronic illness than those in their 80s and 90s (Kheirbek et al. 2017), and longevity is heritable with an estimated heritability around 25%, this suggests that healthy aging is unlikely a random event, but that there are underlying biological mechanisms favorably interplayed with certain environmental factors. A key goal of aging and geroscience research is to reveal these mechanisms to identify effective methods to promote the human healthspan.

Multiple approaches have been explored to achieve this goal. For example, studies have been performed to identify genetic factors associated with longevity which identified a few genes (such as *APOE* and *FOXO3*) that are considered reproducible (Yashin et al. 2018). It is somewhat surprising to see that many disease risk alleles’ occurrences are not significantly different in centenarians versus control population, making it a reasonable hypothesis that some protective factors exist to counter the deleterious effect of the disease risk genes (Bergman et al. 2007). In addition to searching for these protective genetic factors, people have also hypothesized that healthy aging individuals maintain a favorable epigenetic profile and optimal gene expression that are critical for longevity and disease prevention (Brooks-Wilson 2013). However, the exact epigenetic and gene expression profiles defining a healthy aging are still unknown.

Gene expression and other types of “omics” data have been widely used to study the process of aging (Edwards et al. 2007; Glass et al. 2013; Peters et al. 2015; Yang et al. 2015). For examples, epigenome and transcriptome landscapes with aging in mice have revealed widespread induction of inflammatory responses (Benayoun et al. 2019); likewise, the down-regulation of mitochondrial genes across human tissues has been widely reported (Glass et al. 2013; Yang et al. 2015). Studying transcriptomes across multiple species with varied lifespans has similarly reveled a potential role for gene expression regulation in contributing to longer lifespans (Ma et al. 2018). Multiple gene expression based aging hallmarks (Frenk and Houseley 2018) are in concordance with the well-known aging hallmarks summarized by Lopez-Otin et al (2013). In addition to gene expression, epigenetic clock of aging has also been developed based on DNA-methylation markers (Horvath and Raj 2018). These recent studies provide support that gene expression and epigenetic regulations can inform our understanding of human aging.

Despite recent developments in transcriptomic and epigenetic aging research, limited studies have focused on understanding the processes involved in healthy human aging. To investigate how healthy aging is different from aging in the general population at a systems level, one approach is to profile tissues from a healthy aging cohort and compare these to the common aging population, to identify gene regulations specific to the healthy aging. However, obtaining essential tissues from healthy individuals is challenging for ethical and practical reasons. On the other hand, several large-scale human genomic datasets are available and can be repurposed for aging research. Here, we leveraged GTEx data (Consortium et al. 2017) to investigate the difference between “healthy” and “common” aging in humans and study their connection with diseases using a transcriptomic analysis.

We selected GTEx donors who were not annotated with diseases to approximate a “healthy” cohort. We calculated age-associated gene expressions in this “healthy” cohort and compared it to age-associated genes from the “common” cohort which included all GTEx donors. We also compared the “healthy” and “common” aging gene expression with various disease-associated gene expressions to elucidate the relationships among “healthy” aging, “common” aging, and diseases.

## Results

### Data overview

We obtained gene expression and genotype data from the GTEx V7 release and examined 46 tissues with over 80 samples per tissue type, for a total of 11,705 samples, covering 620 donors (Supplemental Table S1, 2, Supplemental Fig. S1).

The GTEx data contained detailed information for each sample and donor, including the donor’s age, post-mortem interval (PMI), RNA integrity number (RIN) and additional information. GTEx donor ages were between 20 to 70 years, and donor PMI was defined as the time between death to the start of sample collection procedure. For each tissue sample, PMI was defined as the time spanning the window from the moment of death (or cessation of blood flow), until tissue stabilization and/or preservation takes place, with values ranging from 29 to 1,739 minutes (Supplemental Fig. S2 & Table S3). We used the sample PMI for majority of the tissues; while we used donor ischemia time for samples from 11 brain subregions, since GTEx did not contain sample ischemic time information for them (see Methods).

### The influence of PMI on age-associated gene identification is tissue-specific

Previous studies reported that some genes’ expression continued to change after death in a tissue-specific manner (Ferreira et al. 2018). As the majority of the GTEx samples were post-mortem samples, some gene expression changes due to PMI could potentially interfere with our age-associated gene expression identification. To ensure that our age-associated gene identification will not be confounded by PMI, we started our analysis by evaluating the relationship between PMI and donor age. We observed a significant positive Pearson correlation between PMI and chronological age in adipose, tibial artery, tibial nerve, skeletal muscle and whole blood (the correlation coefficients were between 0.008 to 0.010, and the Benjamini-Hochberg false discovery rate (FDR) ranged from 2.52×10^−7^ to 2.77×10^−5^). PMI showed no correlation with age in other tissues including all brain subregions, pituitary and terminal ileum, etc. (Supplemental Fig. S3a and Table S4). In contrast, RIN was negatively associated with age in most tissues, particularly in colon, arteries, esophagus and lung (Supplemental Fig. S3b and Table S4).

To examine the influence of PMI on age-associated gene expression changes, we compared two linear regression models. In the first model, we only adjusted gene expression with sex, RIN, three genotype-based principal components (PCs) and gene expression PCs (see methods). In the second model, PMI was included as an extra covariate. For both regression models, we set FDR cut-off to 0.01 to select significant associations between gene expressions and age.

Our results showed that correcting PMI had clear impact on age-associated gene identification in some tissues (Supplemental Fig. S4). Overall, we observed decreased number of age-associated genes in most tissues when we corrected for PMI (Supplemental Table S5). This pattern was particularly apparent in breast mammary, skin, whole blood, lung and adipose tissues. A few exceptions were prostate and minor salivary gland, in which more age-associated genes were found after the PMI correction. In general, for the 46 tissue types we considered, tibial artery showed the largest number of genes associated with age (briefly called as aging genes) (n=8,709), followed by aorta artery (n=5,826), skeletal muscle (n=4,444), nerve tibial (n=3,619) and subcutaneous fat (n=3,812). In contrast, very few aging genes were identified from brain – spinal cord (n=0), brain – substantia nigra (n=0), pituitary (n=1), small intestine-terminal ileum (n=1) and liver (n=7) (Supplemental Table S5). The wide range and tissue specificity of the size of aging genes is consistent with our previous findings (Yang et al. 2015).

### GTEx human aging signatures reasonably recapitulated aging genes from other independent studies

To evaluate whether aging signatures derived from GTEx data were reproducible, we compared them to age-associated gene lists from five independent studies covering the following tissues: brain (Berchtold et al. 2008), skin (Glass et al. 2013), adipose (Glass et al. 2013), blood (Lu et al. 2018), and lung (de Vries et al. 2017). For each tissue type, we calculated the significance of the overlaps between gene lists from published work and our gene sets.

Age-associated gene lists generated from GTEx showed a significant overlap with genes identified from these independent studies. For example, in brain and lung tissues, more than 30% of our genes were found overlapped with other studies, with p-values of 5.81×10^−176^ and 4.32×10^−59^ respectively (Supplemental Table S6), suggesting the aging signatures from GTEx were comparable with the aging signatures from other independent studies.

Interestingly, despite that fewer age-associated genes could be identified by PMI correction in four out of the five tissues, we consistently observed an increase of the overlap between independently identified aging genes and our aging genes in these four tissues, e.g., from 10.53% to 12.70% in blood, 29.30% to 37.04% in lung (Supplemental Table S6), suggesting that the correction of PMI could help increase the reproducibility and robustness of aging signature identification.

### Determining “common” and “healthy” aging signatures

A key goal of aging research is to study mechanisms underlying healthy aging and based on them to identify effective methods to promote human healthspan. However, our understanding of healthy aging is rather limited. Particularly, a clear understanding of healthy aging at the molecular level is yet to be established. For example, it remains to be studied at a systems level how healthy aging is different from common aging, as aging process is usually accompanied with various chronic diseases (Global Burden of Disease Study 2015). One approach to study this is to collect tissue samples from a healthy aging cohort to identify age-associated gene regulations, and compare with general aging population to determine regulations that are unique to the healthy aging. By definition, a truly healthy individual does not die due to chronic diseases. To obtain essential tissues (liver, brain, heart etc.) from these healthy individuals, it is only possible if such donors experiences death due to some other cause, such as accidents, which could not be predicted. Prompt tissue collection can therefore be challenging. On the other hand, evaluating existing human genomic data and repurposing them for aging research represents a convenient alternative, although the apparent limitation is that the original experiment was not ideally designed for studying aging and therefore might not provide measurements on many age-related traits. Since GTEx cohort contained donors annotated with and without chronic diseases, it represents a unique opportunity to allow us to understand the difference between “common” *vs.* “healthy” aging at a transcriptome level. For each donor, GTEx recorded the donor’s medical condition for 21 disease categories, e.g., ischemic heart disease, cerebrovascular disease, chronic respiratory disease, diabetes mellitus type II and hypertension. We mapped these disease categories to their biologically relevant tissues, such as various lung diseases to the lung tissue; diabetes and obesity to the adipose tissue. In addition, we required a tissue to have a relatively large number of aging genes, and the sample size of disease donors to be relatively large (>20% of all tissue samples). Based on these criteria, subcutaneous fat, tibial artery, aorta artery and lung tissues were selected for further study regarding the difference between “healthy” and “common” aging.

Initially, we defined “common” aging signatures as gene expression changes associated with age in the general GTEx cohort, for which we included all the donors regardless of their disease status (“common” cohort). Since this “common” cohort contains individuals annotated with and without chronic diseases, we used them to approximate the general aging population in the society. The “healthy” aging signature was defined as gene expression changes associated with age in the GTEx cohort excluding individuals annotated with tissue-specific diseases (“healthy” cohort). For example, for subcutaneous fat, the “common” aging signatures were calculated from all subcutaneous fat (total n=385), while the “healthy” aging signatures were identified from donors without type 2 diabetes and whose Body Mass Indexes (BMIs) were <30, which resulted in 236 samples (Fig. 1a). In tibial artery (total n=382), the “healthy” cohort (n=292) was obtained from GTEx donors without ischemic heart disease/heart attack/acute coronary syndrome. In lung (total n=379), the “healthy” cohort (n=257) was obtained from donors without chronic respiratory diseases, asthma or pneumonia (details can be seen in Supplemental Table S7). A linear regression model was then applied to identify aging genes in each of the aging cohorts (equation 2).

**Fig. 1.**
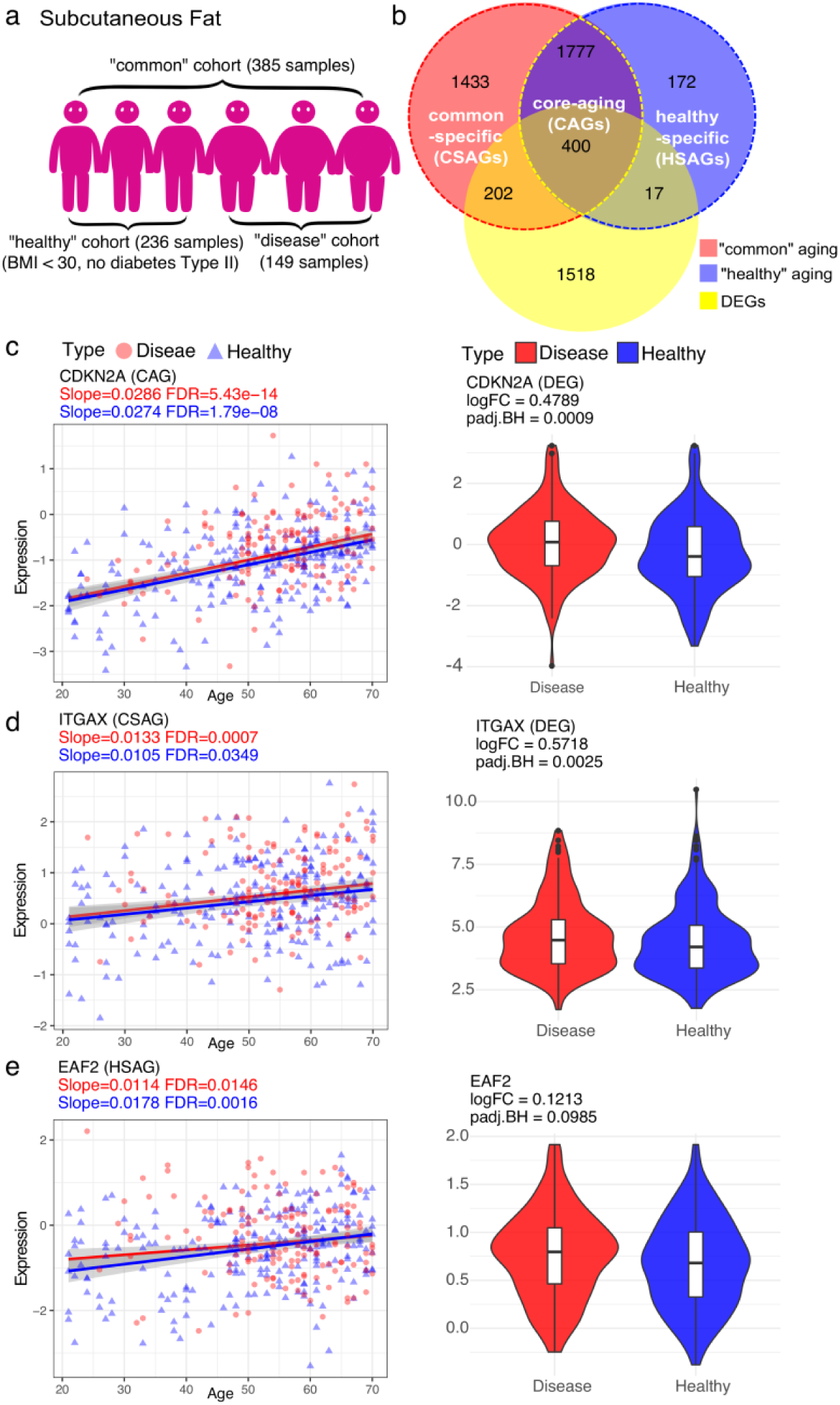
Examples of age-associated gene expression changes in GTEx subcutaneous fat. **a** Cartoon illustration of separating donors into “healthy”, “common” and “disease” cohorts. **b** Venn diagram shows the relationship among “common” aging signature identified from the “common” cohort, “healthy” aging signature from the “healthy” cohort, and DEGs calculated between the “healthy” cohort and the “disease” cohort in subcutaneous fat. **c**,**d**,**e** Scatter plots show 3 representative age-associated gene expression patterns in three gene sets (from top to bottom: “core-aging genes”, “common-specific aging genes”, or “healthy-specific aging genes”). The red line and red dots denote the regression line and samples from the “common” cohort, similarly, the blue line and blue dots represent the regression line and samples from the “healthy” cohort. Violin plots show the gene expression differences between the “healthy” cohort and the “disease” cohort for the corresponding genes in scatter plots. Red violin plot is the gene expression in the “disease” cohort; blue violin plot shows the gene expression in the “healthy” cohort.

It is of note that the “healthy” cohort in this study is more defined at a “healthy” tissue-level rather than a cohort of individuals free of any diseases. For instance, the “healthy” cohort extracted from subcutaneous fat could contain individuals with ischemic heart disease, asthma or chronic obstructive pulmonary disease (COPD). Although theoretically it is ideal to remove individuals annotated with any disease types to obtain a better defined “healthy” cohort, this is not a good option with current GTEx data release since very few samples would be left after this procedure, which could significantly reduce the statistical power of identifying age-associated genes. For instance, only 65 out of 385 samples would remain for subcutaneous fat, and 48 out of 379 samples would remain for lung tissue if we removed all the donors annotated with one or more diseases. Due to this sample size limitation, we chose to approximate our “healthy” cohort in such a tissue-specific way.

### Differentially expressed genes from GTEx “healthy” *vs.* “disease” cohorts were reproducible in other disease-focused transcriptomic studies

As we divided samples based on donor’s disease condition, it is unclear if such division into “healthy” and “common” aging cohorts are biological meaningful. To validate our cohort separation, we calculated the differentially expressed genes (DEGs) between the “healthy” individuals and disease individuals, i.e., we compared 236 “healthy” individuals *vs.* 149 disease individuals in subcutaneous fat, 292 “healthy” individuals *vs.* 90 disease individuals in tibial artery, 198 “healthy” individuals *vs.* 69 disease individuals in aorta artery, and 257 “healthy” individuals *vs.* 122 disease individuals in lung tissue. By using limmavoom (Ritchie et al. 2015), we acquired 2,137 DEGs from subcutaneous fat, 436 DEGs from tibial artery, 1,025 DEGs from lung, while no DEGs were identified from aorta artery. This may be due to the small size of disease samples or other factors such as high heterogeneity in gene expression associated with disease subgroups. We therefore removed aorta artery from the DEG-related analysis (Supplemental Table S7). We then computed the significance of the overlap between our DEGs and disease-related gene signatures obtained from other studies.

Our results showed that DEGs from GTEx significantly overlapped with disease signatures from prior independent studies in the corresponding tissues (Supplementary Table S8). For example, Soronen et al. (Soronen et al. 2012) reported 148 insulin resistance related genes from subcutaneous adipose tissue, 81 of which overlapped with GTEx DEGs in subcutaneous fat, with a p-value of 1.23×10^−42^. These DEGs showed no significant overlap enrichment in artery or lung, suggesting the disease-associated genes were highly tissue-specific. Similarly, genes associated with coronary heart disease (CHD) identified from peripheral whole blood (Joehanes et al. 2013) were exclusively enriched for GTEx tibial artery DEGs (p-value=0.03). Obesity related genes obtained from fat (Font-Clos et al. 2017) were overrepresented only in DEGs from subcutaneous fat and lung (p-values=7.90×10^−4^ and 2.28×10^−3^), but not in tibial artery. Genes involved in chronic obstructive pulmonary disease (COPD) detected from lung (Wang et al. 2008) were found strongly enriched for our GTEx lung DEGs (p-value=5.79×10^−4^), while not in other tissue types. Together, these comparisons indicated that our separation of “healthy” and “disease” individuals based on the GTEx donor health information for the corresponding tissue types was biologically meaningful.

### Characterizing “healthy” and “common” aging signatures suggests some “common” aging genes could be driven by diseases while “healthy” aging signature contained some protective genes

To compare “healthy” and “common” aging signatures, we divided them into three groups: aging genes only seen in the “common” cohort (“common-specific aging genes”, CSAGs), aging genes only observed in the “healthy” cohort (“healthy-specific aging genes”, HSAGs), and common aging genes identified from both cohorts (“core-aging genes”, CAGs). In general, we observed a large overlap between “common” and “healthy” aging signatures in all the four tissues we inspected. For example, in subcutaneous fat, 3,812 “common” aging genes overlapped with 2,366 “healthy” aging genes by 2,177 genes (Fig. 1b and Supplemental Table S7), suggesting that although “healthy” aging is different from “common” aging, they do share a large number of gene expression regulations. In other words, there exists a core aging program regardless the health status of the aging individuals.

We provided examples of gene regulation for CAGs, CSAGs, and HSAGs in Fig. 1. As an example of CSAGs, the expression of integrin subunit alpha X (*ITGAX*) was significantly positively correlated with age in subcutaneous fat in the “common” cohort (FDR=7.12×10^−4^), while its association with age was much less significant in the “healthy” cohort (FDR=0.03) (Fig. 1d). *ITGAX* encodes the integrin alpha X chain protein (also named CD11c), which forms a leukocyte-specific integrin when combined with the beta 2 chain (*ITGB2*). Previous studies reported that CD11c expression in adipose tissue significantly increased in both diet-induced obesity mice and humans (Wu et al. 2010). This is consistent with our results as *ITGAX* was up-regulated in GTEx adipose tissues of donors with high BMIs and type 2 diabetes (Fig. 2d). Since this gene had a positive correlation with age in the “common” cohort but not so when disease individuals were removed, it indicates that its association with age in the “common” cohort was partially driven by the disease individuals. As can be seen in Fig. 1a, over 1,600 CSAGs were identified in the subcutaneous fat, and more than 200 of them were disease DEGs, suggesting that many aging genes derived from the “common” cohort could be partially driven by disease. In contrast, as an example of HSAGs, gene expression changes of ELL-associated factor 2 (*EAF2*) showed strong up-regulation with age (FDR=1.58×10^−3^) in the “healthy” cohort (Fig. 1e), but to a less degree in the “common” cohort (FDR=0.015). *EAF2* gene was found to regulate DNA repair and provide protection to DNA damage in prostate cancer cells (Ai et al. 2017). Lastly, for CAGs, the expression changes of cyclin dependent kinase inhibitor 2A (*CDKN2A*) was substantially associated with age (FDR=5.43×10^−14^ and 1.79×10^−8^) in both cohorts (Fig. 1c). *CDKN2A* encodes for INK4 family member p16 (or p16^INK4a^) which is a well-recognized cell senescence marker (Coppé et al. 2011). The increased expression of *CDKN2A* has been suggested as a biomarker of physiological age (Krishnamurthy et al. 2004).

**Fig. 2.**
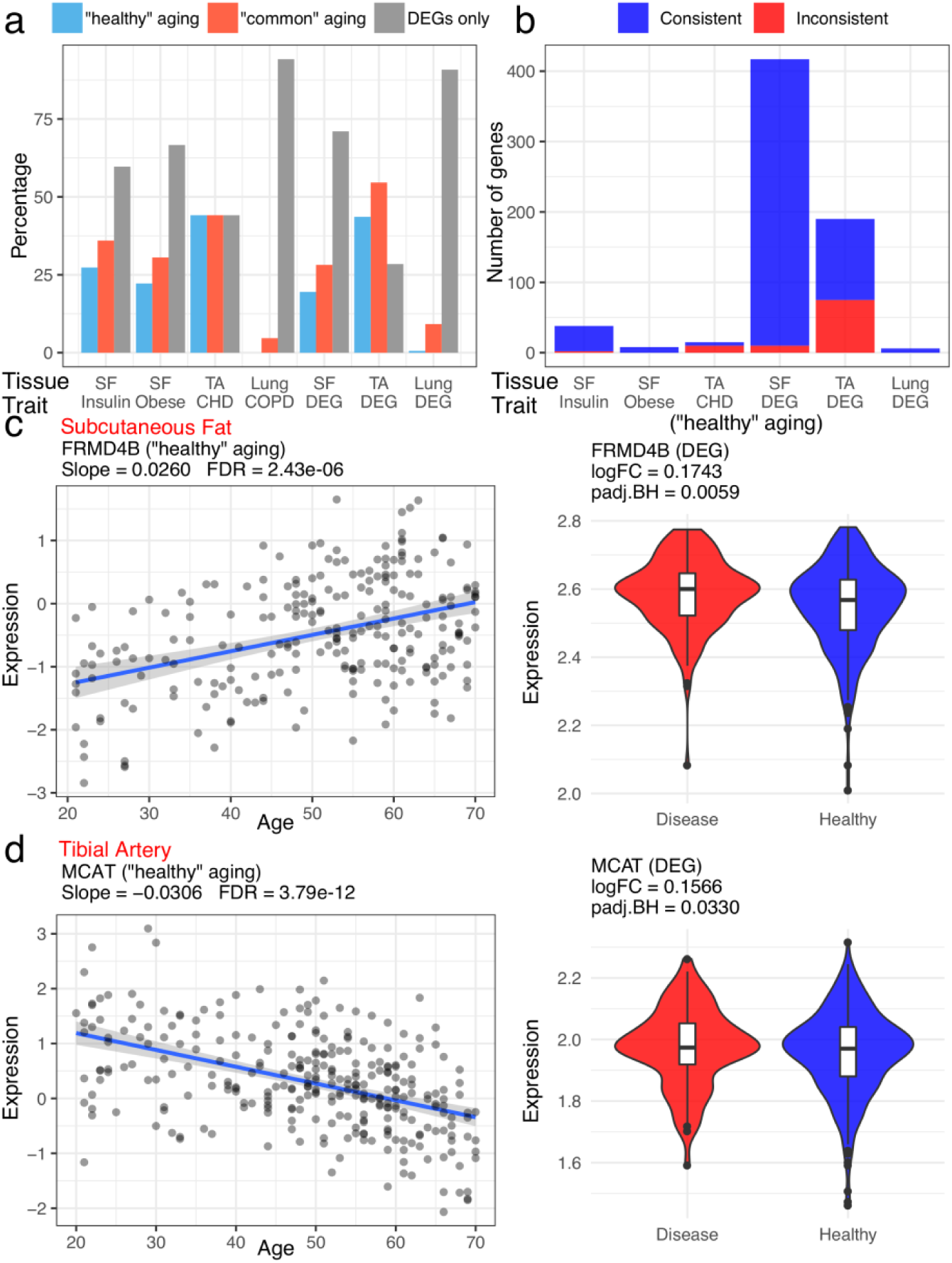
The relationship between age-associated vs. disease-associated gene expression changes. **a.** The percentage of aging signatures (blue: “healthy” aging; red: “common” aging) that overlapped between disease DEGs and the percentage of disease DEGs that were not associated with age (grey bars). Disease DEGs were 4 disease signatures from prior work (insulin, obesity, coronary heart disease (CHD), and chronic obstructive pulmonary disease (COPD) related DEGs) and 3 disease DEGs based on GTEx analysis (simply labeled as DEG). The tissues plotted were subcutaneous fat (SF), tibial artery (TA), and lung. **b** The number of “healthy” aging signatures whose direction was consistent (blue) or inconsistent (red) with the direction of gene regulations in 6 disease DEGs as in **a** (there was no overlapped genes between COPD and “healthy” aging genes in lung). **c**, **d** Two examples of gene expression patterns for “healthy” aging genes. **c**: “Healthy” aging in subcutaneous fat showed the same direction of gene expression change with disease DEG, **d**: “healthy” aging in tibial artery showed an opposite direction of gene expression change with disease DEG. The regression was based on the “healthy” cohort. Violin plots show the gene expression differences between “healthy” (blue) and “disease” (red) individuals.

In addition to having different age-association for their gene expressions, we found that CAGs and CSAGs had significant higher overlap with disease-associated DEGs compared to HSAGs. As shown for subcutaneous fat (Fig. 1b), 400 and 202 genes from CAGs and CSAGs were disease associated DEGs (p-values=5.40×10^−3^ and 1.36×10^−32^, respectively, Supplemental Table S9.). In contrast, only 17 disease DEGs overlapped with HSAGs (p-value=0.78). Similarly, CAGs from tibial artery were specially enriched for its disease DEGs (p-value=9.73×10^−10^), and lung CSAGs were also overrepresented in lung disease associated DEGs (p-value=3.09×10^−11^). While HSAGs were found much less significantly enriched for disease DEGs in tibial artery (p-value=0.03) and lung (p-value=1.00).

### The direction of aging gene expression is largely consistent with the regulation direction in age-related diseases

One key question in geroscience is to understand the underlying mechanisms why aging dramatically increases the incidence of various age-related diseases. Since we had obtained aging signatures from both “healthy” and “common” cohorts, and signatures associated with several diseases, we examined the relationships between “healthy”, “common” aging and disease.

We compared different aging signatures with 7 disease signatures, 4 from previous independent studies (insulin resistance related, obesity related, CHD and COPD) and 3 from DEGs calculated from GTEx data. We observed that most of the disease DEGs were not genes in our aging signatures (Supplemental Table S10). Using insulin resistance/obesity related DEGs in subcutaneous fat as an example, over 60% of them were not associated with age. Furthermore, as previously noticed, “healthy” aging was less enriched for disease DEGs compared with “common” aging. For example, in subcutaneous fat, 36% of “common” aging signatures were found as insulin resistance DEGs, while only 27% of “healthy” aging signatures were insulin resistance DEGs (Fig. 2a).

Although disease and aging signatures were largely different, a substantial number of gene expression regulations were common between them. We considered it interesting to test if age-associated gene expression (particularly in the “healthy” aging) would have similar directions as the regulation changes in disease conditions. We decided to focus on the “healthy” aging since the “common” aging cohort contained disease individuals, therefore the “common” aging genes were not defined independently from disease-associated gene regulations. On the other hand, “healthy” aging genes were identified from the “healthy” cohort, which were considered disjoint from the disease DEGs detected from the “disease” cohort. Our results showed that in most cases, the direction of aging gene regulation in “healthy” aging was largely consistent with the direction of gene expression changes in disease (Fig. 2b&c). Using FERM domain containing 4B (*FRMD4B*) in subcutaneous fat as an example, it was an insulin resistance DEG (also a disease DEG in our GTEx analysis). It is an inflammation-related gene whose expression was up-regulated in the adipose tissue in insulin resistant compared to insulin sensitive group (Wiklund et al. 2016). Our results also showed that the gene expression of *FRMD4B* was up-regulated with increasing age and in disease condition. This consistency suggests that even under “healthy” aging condition, some genes’ regulation is towards a direction that transforms the tissue into a more disease-prone state.

While the direction of gene expression regulation is largely consistent between “healthy” and disease signatures, some gene regulations showed opposite directions of regulation (Fig. 2d). For example, malonyl-CoA-Acyl carrier protein transacylase (*MCAT*), a gene related to pathways like fatty acid metabolism and mitochondrial fatty acid beta-oxidation, its gene expression was found negatively correlated to plasma HDL levels (Ma et al. 2007). In GTEx data, *MCAT* showed a down-regulation in “healthy” aging but its expression was up-regulated in disease population in the tibial artery, suggesting this gene could play a protective role in “healthy” aging to counter the deleterious effect of other disease risk genes.

### Difference in functional enrichment between “healthy” and “common” aging signatures

We investigated the function similarity and difference between “healthy” and “common” aging signatures using DAVID tools (Dennis et al. 2003). Genes were divided into up-regulated *vs.* down-regulated genes with respect to age and were annotated separately (Supplemental Table S11).

GO annotations and pathway analysis revealed differential regulation of genes involved in CAGs, CSAGs and HSAGs. Among genes down-regulated with age, subcutaneous fat CAGs and CSAGs and tibial artery CAGs were characterized with changes in mitochondrial function, energy/oxidation derivation, TCA cycle and genes associated with several neurodegenerative diseases. The cumulative damage to mitochondria and mitochondrial DNA caused by reactive oxygen species is a well-recognized cause of aging (Cui et al. 2012). Down-regulated HSAGs in tibial artery were mainly related to ribosome, various kinds of RNA processing and the regulation of translation; Down-regulated CSAGs in lung were found to be involved in cell-cycle among the top differentially expressed GO terms, while no functional enrichments were found in lung HSAGs (Fig. 3, supplementary Table S12).

**Fig. 3.**
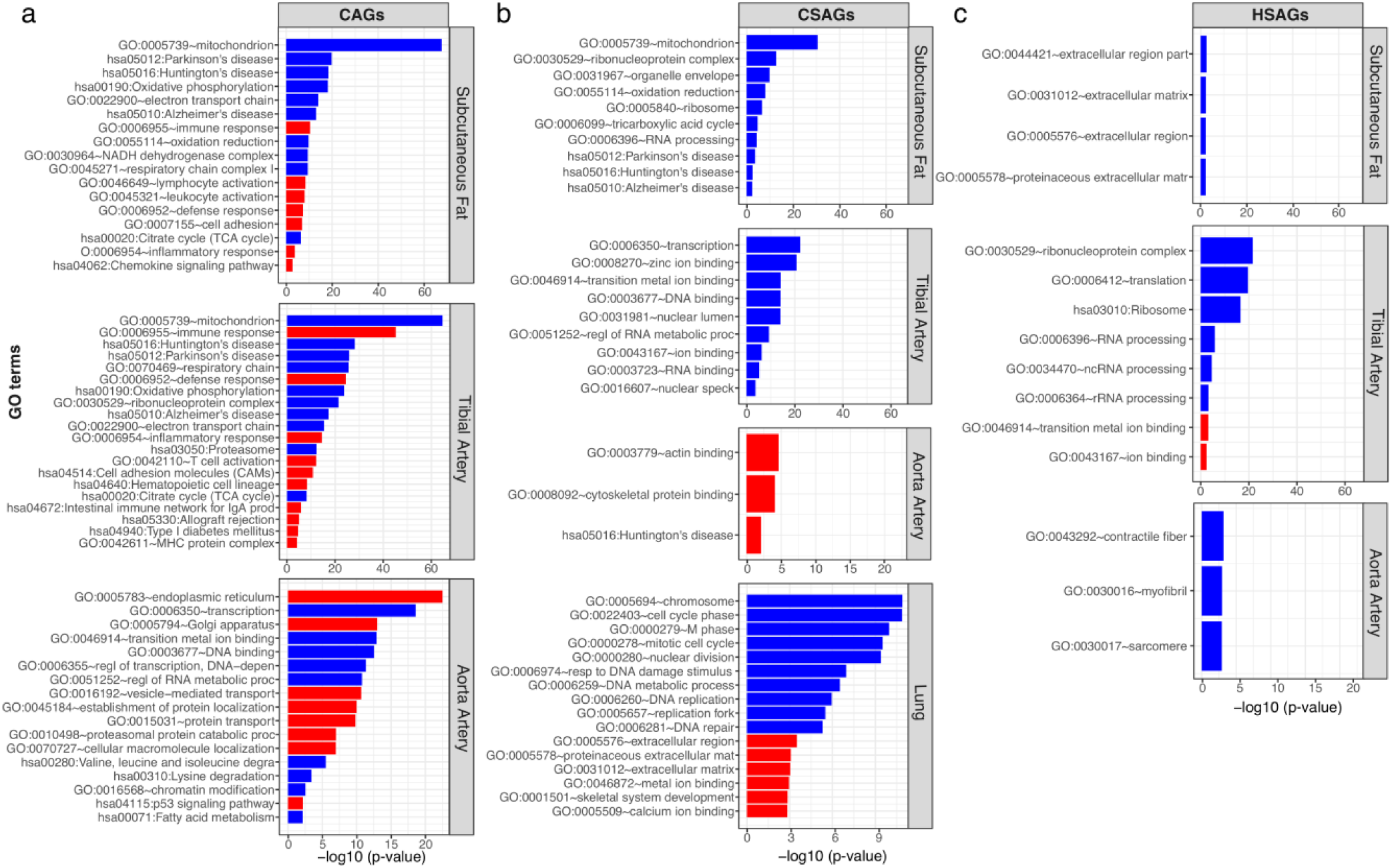
Function enrichment of the “core-aging genes”, “common-specific aging genes” and “healthy-specific aging genes”. **a.** GO terms and KEGG pathways enriched in the CAGs in three tissues (from top to bottom: subcutaneous fat, tibial artery and aorta artery). **b.** GO terms and KEGG pathways enriched in the CSAGs in four tissues (from top to bottom: subcutaneous fat, tibial artery, aorta artery and lung). **c.** GO terms and KEGG pathways enriched in the HSAGs in three tissues (from top to bottom: subcutaneous fat, tibial artery and aorta artery). The red bars denote up-regulated genes with age, the blue bars represent down-regulated genes with age.

Genes up-regulated with age were enriched for a set of different functions compared with genes down-regulated with age (Fig. 3). For example, CAGs in subcutaneous fat were characterized for response to immune/defense/inflammatory, T cell receptor signaling pathway and chemokine signaling pathway, and it was reported that aging is associated with increased T-cell chemokine expression (Chen et al. 2003). Up-regulated CAGs in tibial artery were enriched for similar functions as adipose tissue, in addition, they were also found to be associated with intestinal immune network for IgA production, MHC protein complex and various pathologies. Very few functions were found in up-regulated CSAGs and HSAGs, and they were mostly associated with extracellular component (A full list of functional annotations is provided in Supplemental Table S12).

### Link GTEx age-associated gene expression with known disease and human age genes

Previously, we compiled a list of disease genes for 277 diseases/traits based on the NIH Genome-wide association study (GWAS) catalog (Welter et al. 2014) and Online Mendelian Inheritance in Man (OMIM) (Amberger et al. 2015)(see Methods for details). Using this combined gene list, we investigated the disease gene enrichment for CAGs, CSAGs and HSAGs in four tissues, considering the age-associated up-and down-regulated genes separately. To visualize the results, we displayed the top 10 disease/traits that had significant enrichment in at least one type of age-associated genes for each tissue in Fig. 4. A total of 75 unique diseases/traits with significant overrepresented in down-regulated aging signatures, and 74 unique diseases/traits with significant overrepresented in up-regulated aging signatures were displayed (Fig. 4). We found that disease gene enrichment for “healthy” and “common” aging signatures varied in a tissue-specific manner. For the up-regulated aging genes, most of the significant disease enrichment was observed in either CAGs or CSAGs and to a lesser degree in the HSAGs (Fig. 4a). For example, in tibial artery, CAGs were found to be strongly associated with multiple bowel diseases, including ulcerative colitis, inflammatory bowel disease, crohn’s disease and celiac disease, but very few enrichments were observed for the HSAGs. One of the top up-regulated CAGs in tibial artery, *EDA2R* (FDR=7.67×10^−30^), was considered as a strong candidate gene for lung aging (de Vries et al. 2017).

**Fig. 4.**
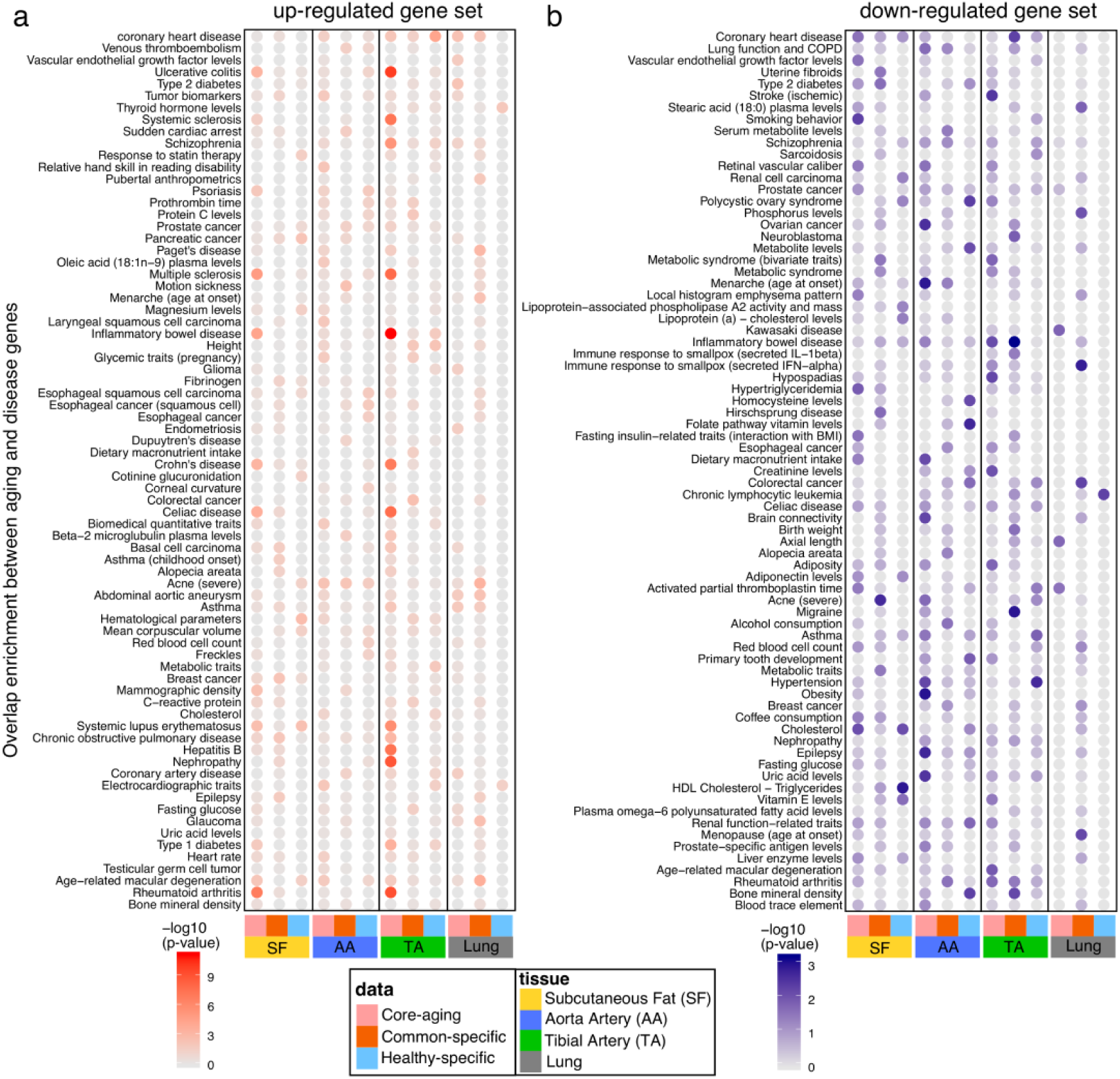
Enrichment of disease genes in the three types of aging gene sets. **A.** The enrichment between up-regulated aging genes and complex disease genes in three aging gene sets (from left to right: CAGs, CSAGs or HSAGs) corresponding to four tissues (subcutaneous fat (SF); aorta artery (AA), tibial artery (TA), and Lung). **b** The enrichment between down-regulated aging genes and complex disease genes in three aging gene sets. Minus log10 transformed p-values were displayer in a color-scale with deeper colors corresponding to more significant p-values.

For genes decreased with chronological age, we generally observed more significant overlaps with various disease traits for all types of aging genes, including the HSAGs (Fig. 4b). For example, in the tibial artery, the HSAGs were significantly enriched for genes associated with hypertension (p-value=1.88×10^−3^); but the enrichment was not observed for the CAGs (p-value=0.82) or CSAGs (p-value=0.57). This is not totally surprising for HSAGs, since down-regulation of such disease-associated gene expression could be beneficial to the healthy aging. Overall, the disease gene enrichment analysis suggests differential regulation of disease genes (fewer up-regulated genes while more down-regulation) with respect to “common” and “healthy” aging.

Last but not least, we tested the enrichment of literature curated candidate human aging genes with respect to “healthy” and “common” aging signatures. 307 candidate human age genes were downloaded from GenAge (de Magalhaes and Toussaint 2004). We then calculated the overlap enrichment with three aging gene sets (Table 1). Our results showed that CAGs in subcutaneous fat and aorta artery were enriched for human aging genes from GenAge (p-value < 0.01), while neither the CSAGs or HSAGs were enriched for them.

**Table 1.** Comparison of three aging signatures with human aging genes from GenAge. We compared “core-aging genes”, “common-specific aging genes” and “healthy-specific aging genes” from subcutaneous fat, aorta artery, tibial artery and lung with human aging genes from GenAge. No. of background genes was set as the number of all genes we used to calculate age-associated genes in each tissue; No. of GenAge contains the number of overlapped genes between GenAge and our background genes *vs.* the total number of original GenAge genes; p-value was calculated by Fisher’s exact test.

| Tissue |  | GenAge |  |  |
| --- | --- | --- | --- | --- |
|  |  | Core-aging | Common-specific | Healthy-specific |
| Subcutaneous Fat | No. of background genes | 18,643 |  |  |
|  | No. of GenAge genes | 282/307 |  |  |
|  | No. of aging genes | 2,177 | 1,635 | 189 |
|  | No. of overlapped genes | 54 | 24 | 3 |
|  | p-value | 0.0002 | 0.5927 | 0.5470 |
| Aorta Artery | No. of background genes | 18,189 |  |  |
|  | No. of GenAge genes | 281/307 |  |  |
|  | No. of aging genes | 4,321 | 1,505 | 389 |
|  | No. of overlapped genes | 91 | 24 | 10 |
|  | p-value | 0.0006 | 0.4666 | 0.0807 |
| Tibial Artery | No. of background genes | 17,708 |  |  |
|  | No. of GenAge genes | 278/307 |  |  |
|  | No. of aging genes | 5,488 | 3,221 | 2,686 |
|  | No. of overlapped genes | 92 | 53 | 28 |
|  | p-value | 0.2412 | 0.3754 | 0.9952 |
| Lung | No. of background genes | 19,540 |  |  |
|  | No. of GenAge genes | 285/307 |  |  |
|  | No. of aging genes | 96 | 884 | 11 |
|  | No. of overlapped genes | 1 | 22 | 0 |
|  | p-value | 0.7568 | 0.0106 | 1.0000 |

## Discussion

We leveraged GTEx data to study the difference between “healthy” and “common” aging at a transcriptome level. We constructed two cohorts: one containing individuals without diseases related to the tissue type under investigation and one containing all individuals regardless of their disease status. We then identified and compared age-associated gene expressions patterns in these cohorts and also compared them with disease-associated gene expression.

Our work suggests an intimate intertwined relationship between aging and disease in their gene regulations. We have shown that “healthy” and “common” aging shared a large proportion of genes, suggesting the existence of a core aging program regardless of the individual’s health status. This is not surprising, since our “healthy” cohort was a subset of the “common” aging cohort. For genes whose expression in disease individuals are not dramatically different from the “healthy” aging cohort (particularly for the direction of gene expression regulation), adding these disease individuals to the “healthy” individuals will unlikely change the overall gene expression pattern. In addition, we found our core aging genes were enriched for candidate human age genes recorded in GenAge. Therefore, this core aging program likely reflects the main aging regulation in humans since it is insensitive to the health status of the individuals enrolled for a particular aging study.

Despite the large overlap between “healthy” and “common” aging genes, HSAGs and CSAGs showed different function enrichment, and CAGs/CSAGs showed higher enrichment for disease genes. For example, CAGs and CSAGs in subcutaneous fat and tibial artery were enriched for genes related to neurodegenerative disease (Supplemental Table S12). It is of note that aging genes in adipose and artery have been reported to be associated with several chronic neurodegenerative diseases (Balistreri et al. 2010; Akinyemi et al. 2013; Parimisetty et al. 2016). Since certain CSAGs become age-associated only when disease individuals are included, their association with age in the “common” cohort is therefore likely driven by diseases.

Disease associated gene expression regulations are overall different from age-associated genes, supporting that aging and disease are fundamentally distinct and depend on different gene regulations. However, disease associated transcriptome signatures do share some common genes with “healthy” aging signatures. For these shared genes, the direction of gene regulation in “healthy” aging is largely consistent with the regulation direction induced by disease. This suggests that transcriptome regulation in healthy aging could facilitate the development of disease. For example, even in the “healthy” aging adipose tissues, we observed elevated inflammation gene expression (e.g., *CDKN2A, IL4R, TGFB1* and *PTPN22* expression in CAGs, and *TNFS4F* expression in the HSAGs), and it has been noticed that obesity-associated chronic low-grade inflammation is responsible for the decrease of insulin sensitivity (Chen et al. 2015), suggesting that age-associated gene regulation could facilitate the development of certain age-related diseases.

In contrast to the “common” aging signatures which were enriched for disease genes, HSAGs (particularly the up-regulated HSAGs) were less enriched for disease genes and showed different associated functions. For example, down-regulated CAGs in subcutaneous fat were overrepresented for disease traits like cholesterol and hypertriglyceridemia; while down-regulated HSAGs were strongly related to HDL cholesterol-triglycerides (“good” cholesterol). HSAGs also showed strong enrichment for genes related to ribosome and RNA processing; previous studies have reported that both caloric restriction and rapamycin treatment extend health/life-span but substantially decrease mRNA levels of ribosomal proteins through reduced mTOR activity (Frenk and Houseley 2018), therefore that HSAGs may regulate the ribosomal proteins for the benefit of healthy tissue aging.

We speculated that some “healthy-specific” aging genes provide protective mechanisms to prevent the development of diseases. Using adipose tissue as an example, we found that the top few up-regulated genes were *KLF4* and *EAF2* (Supplemental S2_Data). The *EAF2* gene (ELL-associated factor 2) has complex and overall protective functions in different cell and tissue types. For example, *EAF2* is a key factor mediating androgen protection of DNA damage via Ku70/Ku80 in prostate cancer cells (Consortium et al. 2017). It may also suppress oxidative stress-induced apoptosis of HLE-B3 cells exerted through the activation of Wnt3a signaling (Feng and Guo 2018) and protect cells against hypoxia-induced cell death and inhibit cellular uptake of glucose under hypoxic conditions (Chen et al. 2014). *KLF4* functions as an immediate-early regulator of adipogenesis specifically induced in response to cAMP (Birsoy et al. 2008), while abiogenesis is known to be reduced in elderly individuals and correlates with the deteriorated functions of old adipose tissues (Kirkland et al. 2002). It could be an important feature for the healthy aging program to up-regulate multiple protective genes to strengthen the resilience in these aging tissues.

We pointed out that one limitation of our study resides in how we defined a “healthy” aging cohort. It may be argued that our definition of the “healthy” aging cohort is not truly healthy and better criteria should be used. Although we fully understand and agree on this point, we are limited by the available samples in the current GTEx release. When the full version GTEx data become available in near future, we expect to have a sample size that we can define a healthier “healthy” aging cohort. On the other hand, it has been and will continue to be difficult to collect tissues from a truly healthy living person as aging is normally accompanied by various kinds of diseases and only less than 20% of older individuals are considered really healthy. Previous studies focused on “healthy” aging used peripheral blood samples from a living cohort (Lunnon et al. 2012; Erikson et al. 2016), or muscle samples from people with good aerobic fitness and free from metabolic and cardiovascular disease (Sood et al. 2015), or brain regions that were neurologically normal, but were obtained from donors who died from ischemic heart disease (Ramasamy et al. 2014). Although specific consideration was taken, most of these studies did not require that samples were from absolutely healthy individuals. Therefore, we consider it is acceptable to define the “healthy” cohort as donors without tissue-specific diseases given the resources available to us now, particularly as meaningful results seem to be obtainable by this approach.

In conclusion, we performed a comparative analysis of “healthy” and “common” aging genes based on transcriptomic data from GTEx. We found “common” aging signatures are comparably more associated with genes and pathways that cause disorders during aging process, while “healthy” aging is likely to contain more “good” factors and pathways that engage in boosting the systems resilience in humans. As a future direction, a useful effort would be to catalog the protective aging gene expression regulations in details and discover potential interventions to promote the healthy aging program in “common” aging populations.

## Methods

### Data availability and filters

RNA-seq data (Illumina paired-end, 76bp) and genotyping data from GTEx release V7 were downloaded from the Database of Genotypes and Phenotypes (dbGaP) under accession pht002742.v2.p1 (see Supplemental Table S1). Details about the RNA-seq data are provided in the Supplemental Table S2.

Form the 54 available tissues in the V7, we started by selecting those with at least 80 samples, and samples with more than 20 million mapping reads and greater than 40% mapping rate. Cell line data were removed from our analysis, including EBV-transformed lymphocytes, Leukemia cell line (CML), and transformed fibroblasts. Only genes with expression > 0.1 TPM and aligned read count of 5 or more in more than 80% of all samples within each tissue were considered significantly expressed and used for aging signature identification. Expression measurements for each gene in each tissue were subsequently inverse quantile normalized to a standard normal distribution.

Our final dataset included samples from 46 tissue types. The sample size in each tissue ranged from 85 to 491, with an average of 247 samples. The number of genes varied from 14,441 in whole blood to 27,280 in testis, with a mean of 18,812 genes (Supplemental Fig. S1).

### Age and post-mortem interval (PMI) information

Based on GTEx protocol, only donors within 20-70yrs were collected for their project (Supplemental Fig. S2a). GTEx annotation provides information on three types of ischemic time: Total Ischemic time for a sample (SMTSISCH), Total Ischemic time for a donor (TRDNISCH) and Ischemic Time (TRISCHD) (time for the start of the GTEx procedure). Tissue collection was completed within 24hrs of cardiac cessation. Throughout the text we have used the term Post-Mortem Interval (PMI) to refer to sample ischemic time. For 13 brain subregions provided by GTEx, only brain cerebellum and brain cortex have sample ischemic time information. In those cases, we used donor ischemic time for the rest of the brain subregions, and they are marked with specific notations. Negative PMI values observed in whole-blood samples were transformed to 0 in the regression analysis (Supplemental Fig. S2b,c).

### PMI correction

In order to assess the impact of PMI on age-associated gene expression changes, regression models with or without PMI as confounding factors were applied. First, we considered a set of covariates (age, gender, top 3 genotypes, gene expression PC and RIN). Then, for each gene we implemented a linear regression model where the gene expression was modeled with relation to the covariates (equation 1). Next, the correlation between the residuals and the age was calculated, r = correlation (gene_expression.resid, age.vals). The corresponding correlation and p-values (adjusted with BH (Benjamini and Hochberg 1995) method) were then calculated for all genes; only FDR value < 0.01 were considered as age-associated genes. This procedure was repeated, except by correcting PMI as an extra confounding variate (equation 2).

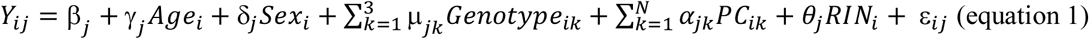

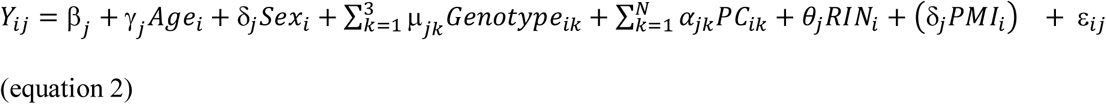

In both models, *Y*_*ij*_ is the expression level of gene j in sample i, *Age*_*i*_ denotes the donor age of sample i, *Sex*_*i*_ denotes the sex of donor for sample i, *Genotype*_*ik*_(*k* ∈ (1,2,3)) denotes the value of k-th principal component value of genotype profile for the i-th sample, *PC*_*ik*_ (*k* ∈ (1, …, *N*) denotes the value of k-th principal component value of gene expression profile for the i-th sample, N is the total number of top PCs under consideration, *RIN*_*i*_ denotes the RIN score of sample i, *PMI*_*i*_ denotes the PMI of sample i, which was considered as an extra covariates in the second regression model (equation 2), *ε*_*ij*_ is the error term, *γ*_*j*_, *δ*_*j*_, *μ*_*jk*_, *δ*_*j*_, is the regression coefficient for each variates. For each gene j, a least square approach was used to estimate the regression coefficients. If *γ*_*j*_ was significantly deviated from 0, gene j was then considered to be an age-associated gene. Gene j with *γ*_*j*_> 0 was considered as up-regulated with age, gene j with *γ*_*j*_< 0 was considered as down-regulated with age.

### Differential expression between the “disease” and “healthy” individuals

For differentially expression analysis we used the statistical methods implemented in the limma-voom (Ritchie et al. 2015) package. We started by building a matrix with gene read counts in donors with tissue-specific diseases and donors without tissue-specific diseases (Supplemental Table S7). Genes are kept if they are expressed in at least two samples. We created a design matrix taking into account 2 conditions (e.g. “disease” cohort and “healthy” cohort) and several covariates:

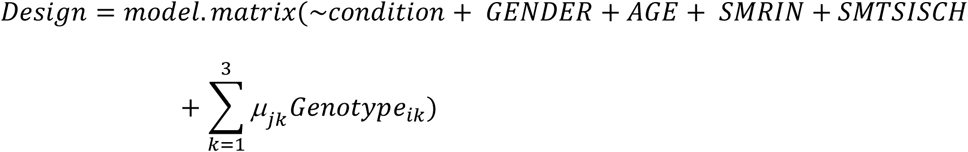

Condition and gender were converted as factors, type of nucleic acid isolation batch (SMNABTCHT) was implemented as batch. The corresponding correlation and p-values (adjusted with BH) were then calculated for all genes, only results with FDR value < 0.05 were considered as significant DEGs.

### Functional annotation for GTEx aging signatures

To demonstrate the potential functional significance of aging signatures, DAVID tool (Dennis et al. 2003) was used to perform GO classification. We first extracted gene symbols from each age-associated gene sets: “common-specific aging genes, “core-aging genes” and “healthy-specific aging genes. These gene lists were then submitted to DAVID by choosing GO_FAT and KEGG pathway terms to describe the overrepresented functional terms for them. The threshold for overrepresented GO terms was set to FDR less than 0.05.

### Assembly of disease gene list and identify significant overlap between disease and aging genes

Disease genes were retrieved from two sources: NIH Genome-Wide Association Studies (GWAS) Catalog and OMIM (Online Mendelian Inheritance in Man). We only considered genes in the GWAS catalog with p-value < 5×10^−8^, a widely accepted threshold for genome-wide significance. Clustering and manual curation were used to merge genes in GWAS and OMIM. We only considered disease categories which contained with at least 5 genes. We then performed a Hypergeometric based test between the disease genes and three age-associated gene sets in four tissues. Aging genes with FDR ≤ 0.01 were used for testing age-disease overlap enrichment (p-values less than 0.01 were considered significant). In addition, we separated each gene set into up and down-regulated with age. To visualize the result, we selected the top 10 most significant diseases in each tissue, which resulted in 75 unique diseases for down-regulated and 74 unique diseases for up-regulated age-associated genes. P-value was then -log10 transformed and plotted in Fig. 4.

### Data access

The GTEx analysis V7 genotype, phenotype and gene expression data were downloaded from GTEx portal: https://gtexportal.org/home/, and dbGaP accession phs000424.v7.p2.

## Acknowledgments

This work was funded by NIH grant R01AG055501 to Z.T. and Y.S. We wish to acknowledge all GTEx donors for their generous tissue donation and GTEx consortium for data curation. This work was supported in part through the computational resources and staff expertise provided by Scientific Computing at the Icahn School of Medicine at Mount Sinai.

## Author contributions

Z.T. conceived and designed the project; L.Z. performed the analysis; L.Z. and Z.T. wrote the paper; J.Y., S.P., J.Z., B.Z., and Y.S. contributed to the discussion of the results and helped revise the paper. All authors reviewed the work and agreed for its publication.

